# Phenylhydrazones are active against non-replicating *Mycobacterium tuberculosis*

**DOI:** 10.1101/323220

**Authors:** Shilah A. Bonnett, Devon Dennison, Anumita Bajpai, Megan Files, Tanya Parish

## Abstract

There is an urgent need for the development of shorter, simpler and more tolerable drugs to treat antibiotic tolerant populations of *Mycobacterium tuberculosis*. We previously identified a series of phenylhydrazones (PHY) active against *M. tuberculosis*. We selected six representative compounds for further analysis. All compounds were active against non-replicating *M. tuberculosis*, with two compounds demonstrating greater activity under hypoxic conditions than aerobic culture. Compounds had bactericidal activity with MBC/MIC of <4 and demonstrated an inoculum-dependent effect against aerobically replicating bacteria. Bacterial kill kinetics demonstrated a faster rate of kill against non-replicating bacilli generated by nutrient starvation. Compounds had limited activity against other bacterial species. In conclusion, we have demonstrated that the PHY compounds have some attractive properties in terms of their anti-tubercular activity.

## Introduction

Tuberculosis (TB), caused by *Mycobacterium tuberculosis*, is a global health problem [1]. In 2016, 10.3 million people worldwide became ill with TB and 1.7 million people lost their lives to the disease [1]. While the number of deaths fell ~ 24%, the number of new cases increased slightly to 6.3 million in 2016. Approximately a quarter of the world’s population has latent TB in which patients are asymptomatic and non-infectious. Reactivation of latent infection is observed in 10% of cases representing a large reservoir of infection [2, 3].

During latent infection, *M. tuberculosis* bacilli can persist in the granuloma for years. During this time, the bacteria are in a slow or non-replicating state with low metabolic activity. The metabolic state of the bacilli is influenced by host environmental conditions such as low oxygen and pH, nutrient deprivation, and exposure to RNS and ROS [4], all of which may contribute to antibiotic tolerance. *In vitro*, starvation-induced, non-replicating bacilli are tolerant to isoniazid, rifampicin and metronidazole, but not pyrazinamide, econazole or clotrimazole [5––13].

The prevalence of latent TB has complicated our ability to eradicate the disease. There is an urgent need for the development of shorter, simpler and more tolerable drug regimens to treat various subpopulations of *M. tuberculosis*. In order to attain a shorter therapy period, new drugs should be bactericidal and be efficacious against non-replicating and antibiotic tolerant forms of *M. tuberculosis*.

The hydrazone linker (-NH-N=CH-) is a useful synthetic tool enabling the generation of hydrazide-hydrazone derivatives, many of which are pharmacologically-active. Such molecules target wide range of diseases and have anti-microbial, anti-cancer, anti-malarial and anti-inflammatory activities [–29]. While the mode of action of hydrazones varies depending upon the structural characteristics, several are involved in covalent modification of proteins and/or sequestering of metal ions.

Isoniazid (INH), a first line TB drug is a hydrazine, which is a tight-binding inhibitor of enoyl reductase (InhA) in *M. tuberculosis*; INH is a prodrug which requires activation by the KatG catalase-peroxidase [––35]. More recently, 2-hydroxy-1-naphthaldehyde isonicotinoyl hydrazone, was identified as a selective inhibitor of *M. tuberculosis* methionine aminopeptidase (MetAPs) with activity against replicating and non-replicating bacteria [36]. In addition, quinolone hydrazone derivatives are currently being explored as potential anti-cancer and anti-tubercular drugs [37]. Interestingly, copper (II) and zinc (II) complexes of quinolone hydrazone derivatives have higher anti-tubercular activity than the free hydrazone [37].

We previously identified a series of phenylhydrazones (PHY) in a target-based whole-cell screen [14]. We demonstrated good activity in growth inhibition assays against actively growing wild-type bacteria, as well as improved activity against a strain engineered to under-express the sole signal peptidase, LepB [14]. We conducted a small structure-activity relationship study and identified several modifications which improved potency against *M. tuberculosis.*

## Materials and Methods

### Bacterial culture

Mycobacteria were cultured in Middlebrook 7H9 medium supplemented with 0.5% w/v Tween 80 and 10% v/v oleic acid, albumin, dextrose, catalase (OADC) supplement (7H9-Tw-OADC) or on Middlebrook 7H10 agar plus 10% v/v OADC. *Escherichia coli* DH5α and *Staphylococcus aureus* RN4220 were grown in LB broth and on LB agar. *Pseudomonas aeruginosa* HER1018 (PAO1) was grown in tryptic soy broth and on tryptic soy agar. *Bacillus subtilis* Marburg was grown in nutrient broth and on nutrient agar. *Saccharomyces cerevisiae* Y187 was grown in YPD broth and on YPD agar supplemented with 0.2 % adenine hemisulfate.

### Minimum inhibitory concentration (MIC) determination

MICs were determined in liquid medium in 96-well, black, clear-bottom plates as described [38]. A 10-point 2-fold serial dilution was run for each compound and bacterial growth was measured by OD_590_ after 5 days of incubation at 37°C. Growth inhibition curves were fitted using the Levenberg–Marquardt algorithm. The IC_90_ was defined as the concentration of compound required to inhibit growth by 90%.

### Low Oxygen Recovery Assay (LORA)

The Low Oxygen Recovery Assay was carried out as described in 96-well plates [5]. Bacteria (*M. tuberculosis* strain H37Rv-LUX) were cultured in Dubos medium with supplement (DTA) in the Wayne Model of hypoxia for 18 days to enter hypoxia and used to seed 96-well plates containing compounds. Plates were incubated for 9 days under anaerobic conditions followed by 28h outgrowth under aerobic conditions; as a comparator plates were incubated for 6 days under aerobic conditions. Growth was measured by luminescence. Growth inhibition curves were fitted using the Levenberg– Marquardt algorithm. IC_90_ was determined as the minimum concentration required to prevent 90% growth.

### Minimum Bactericidal Concentration (MBC)

MBCs were determined as described [39]. Briefly, a late log phase culture (OD_590_ 0.6-1.0) was adjusted to an OD of 0.1 in 7H9-Tw-OADC and 50 μL used to inoculate 5 mL of 7H9-Tw-OADC containing compound. Cultures were incubated standing at 37°C. Bacterial viability was determined by plating serial dilutions and enumerating CFUs after four weeks of incubation at 37°C. For starvation, *M. tuberculosis* H37Rv was resuspended in phosphate buffer saline (PBS) plus 0.05% w/v Tyloxapol and incubate standing for 2 weeks. Compound was added after two weeks and CFUs determined.

### Spectrum

MICs were determined using the serial dilution agar method. Unless otherwise stated, compounds were prepared as an 8-point 2-fold serial dilution in DMSO starting at 100 μM. MIC_99_ was defined as the minimum concentration that prevented 99% of growth.

## Results and discussion

The PHY series have good activity against aerobically-cultured, actively growing *M. tuberculosis* in axenic culture. Our previous work was limited to determining minimum inhibitory concentrations (MICs) under these conditions and demonstrated activity for a range of analogs, with many MICs in the range of 20 μM [14]. However, many compounds that act against rapidly growing mycobacteria are ineffective against non-replicating or intracellular organisms. Since *M. tuberculosis* can survive under low oxygen tension, we were interested to determine whether our compounds had activity under this setting, which is relevant to the environment encountered during infection. We were also interested in determining if compounds were bactericidal against replicating and non-replicating bacilli. We selected six representative PHY compounds based on their activity and structure for characterization in other assay systems (Fig 1) [14].

**Fig 1.**
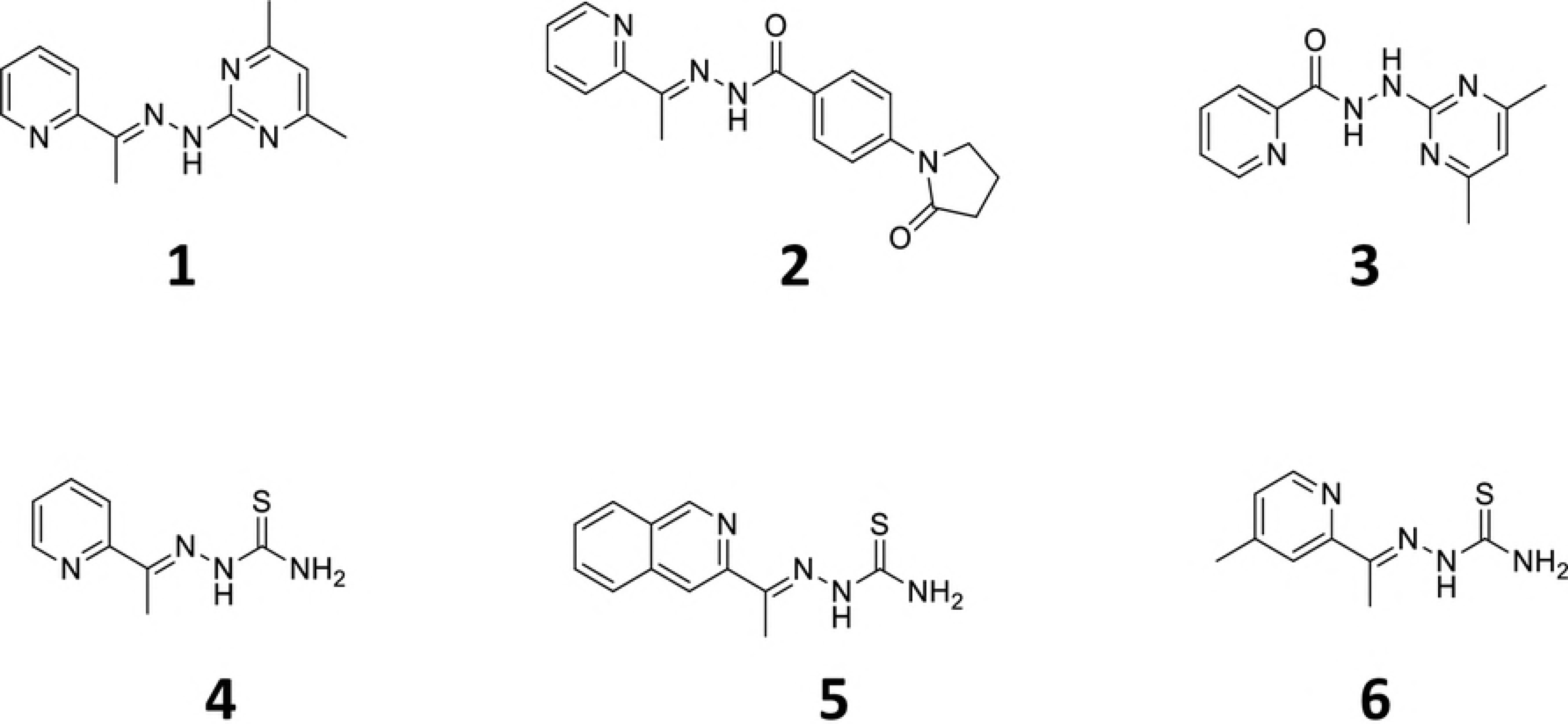
Structure of PHY analogs used in this study.

### PHY compounds are active against hypoxically-induced, non-replicating *M. tuberculosis*

We determined the activity of our compounds using the low-oxygen-recovery assay (LORA) [5, 40]. We determined the IC_90_ (the concentration required to prevent outgrowth by 90%) for bacteria under both aerobic and anaerobic conditions for comparison. Six compounds were tested (Table 1). Five of the compounds were active under aerobic conditions, with IC_90_ < 20 μM. All of the compounds were active under anaerobic conditions, with two compounds, the thiosemicarbazones **4** and **6**, showing greater activity under hypoxia (>2-fold difference). Two compounds (**3** and **5**) were equally active under both conditions. These data demonstrate that the PHY series is efficacious against hypoxia-induced non-replicating bacilli.

**Table 1.** PHY compounds are active under hypoxia. *M. tuberculosis* was cultured in DTA medium in the Wayne model of hypoxia for 18 days to generate the non-replicating state. Bacteria were inoculated into plates containing compounds and incubated for 9 days under hypoxia (anaerobic) or 6 days in air (aerobic). Growth was measured by luminescence. IC_90_ is the concentration required to inhibit growth by 90%. Results are the mean ± standard deviation from a minimum of two experiments.

| Cpd # | Anaerobic IC<sub>90</sub><br>(μM) | Aerobic IC<sub>90</sub><br>(μM) |
| --- | --- | --- |
| 1 | 17 ± 3.0 | > 20 |
| 2 | 22 ± 12 | 6.4 ± 2.4 |
| 3 | 8.3 ± 0.3 | 6.2 ± 4 |
| 4 | 5.5 ± 2.5 | 13 ± 6.3 |
| 5 | 2.5 ± 1.3 | 2.7 |
| 6 | 9.6 ± 0.4 | 20 ± 7 |

### PHY compounds are bactericidal against replicating *M. tuberculosis*

We first determined whether compounds had bactericidal or bacteriostatic activity under aerobic conditions (Fig 2). We tested four compounds at varying concentrations. The highest concentration of compound that can be tested in our assay and keep DMSO to 2% is 200 μM. Therefore we selected concentrations to test as follows: for compounds with MIC <20 μM, we used concentrations as a multiple of the MIC i.e. 10X, 5X, 2.5X, and 1X MIC; for compounds with MICs >20 μM, we used fixed concentrations of 200,100, 50, 25 and 20 μM.

**Fig 2.**
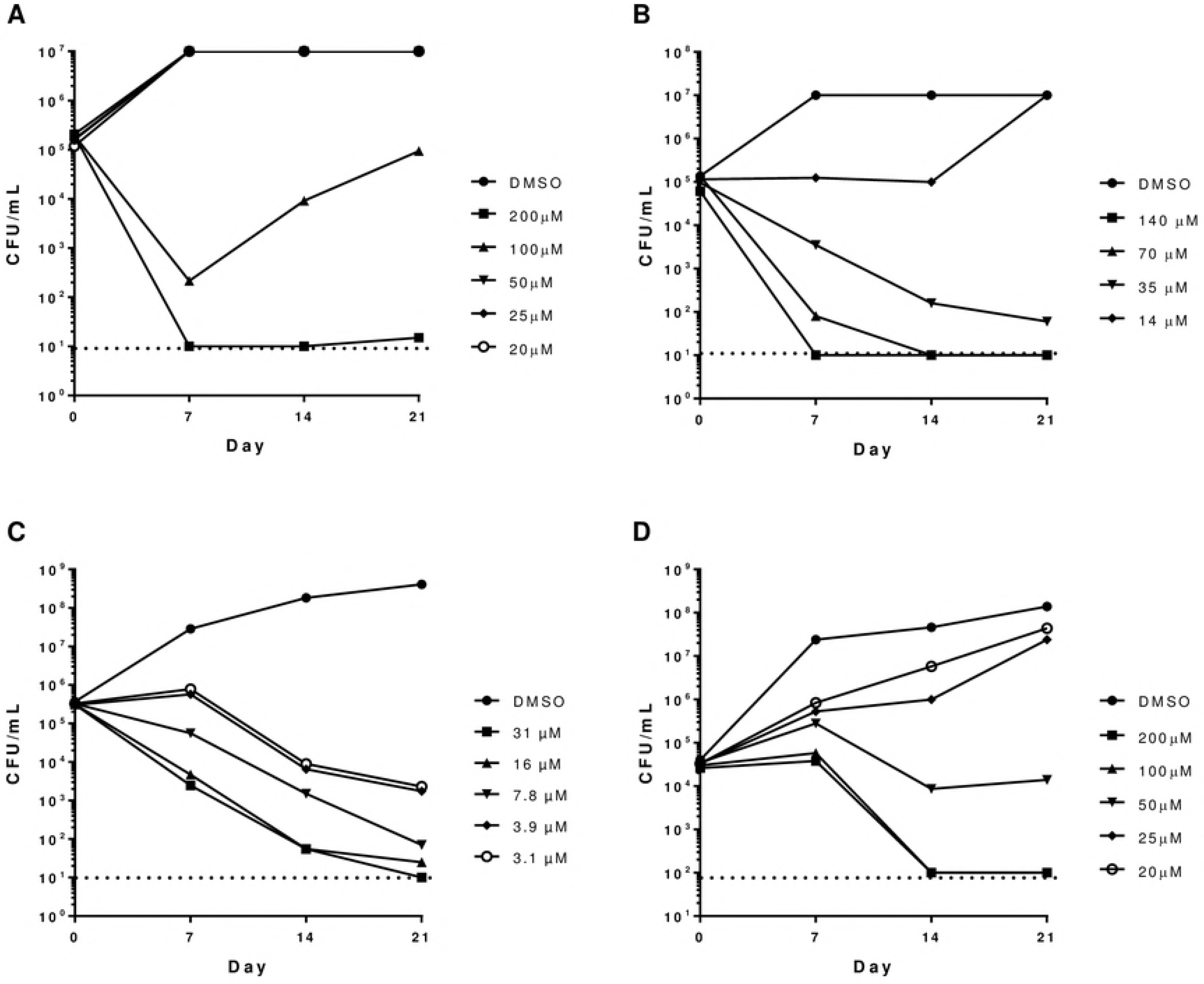
Kill kinetics against replicating *M. tuberculosis*. Bacterial viability in the presence of compound was determined by counting CFU every 7 days over a 21-day period. Compounds (A) **1,** (B) **2**, (C) **3** and (D) **6**. The dashed line represents the lower limit of detection.

All compounds were able to effect a 3 log kill within 21 days. At the highest concentrations compounds **1** and **2** were able to sterilize culture within 7 days, compound **3** within 14 days, and compound **6** within 21 days. Compound activity was concentration-dependent, (as defined by CLSI guidelines [41]), since the rate of kill increased with increasing concentration. The MBC, defined as a 3 log kill within 21 days, was determined; compound **1** was 200 μM, compound **2** was 35 μM, compound **3** was 7.8 μM, and compound **4** was 100 μM. The compounds were all classified as bactericidal i.e. MBC/MIC of <4 [41]. For compound **1**, the increase in CFUs in the culture treated with 100 μM after day 7 is likely due to the outgrowth of resistant mutants at the lower concentration.

### The bactericidal activity of PHY compounds is inoculum-dependent

We examined the effect of inoculum size on the efficacy of compound **1** and **2** against *M. tuberculosis* under aerobic conditions (Fig 3). Cultures were exposed to 10X MIC over a 7 day period. Both compounds behaved in a similar fashion and their effect was inoculum-dependent i.e at high starting inoculum (~10^7^ CFU/mL), compounds had no impact on bacterial viability. At lower inoculum size, a 2 log reduction in CFU/mL was observed. Complete kill by day 7 was only seen when the inoculum was ≤ 10^5^.

**Fig 3.**
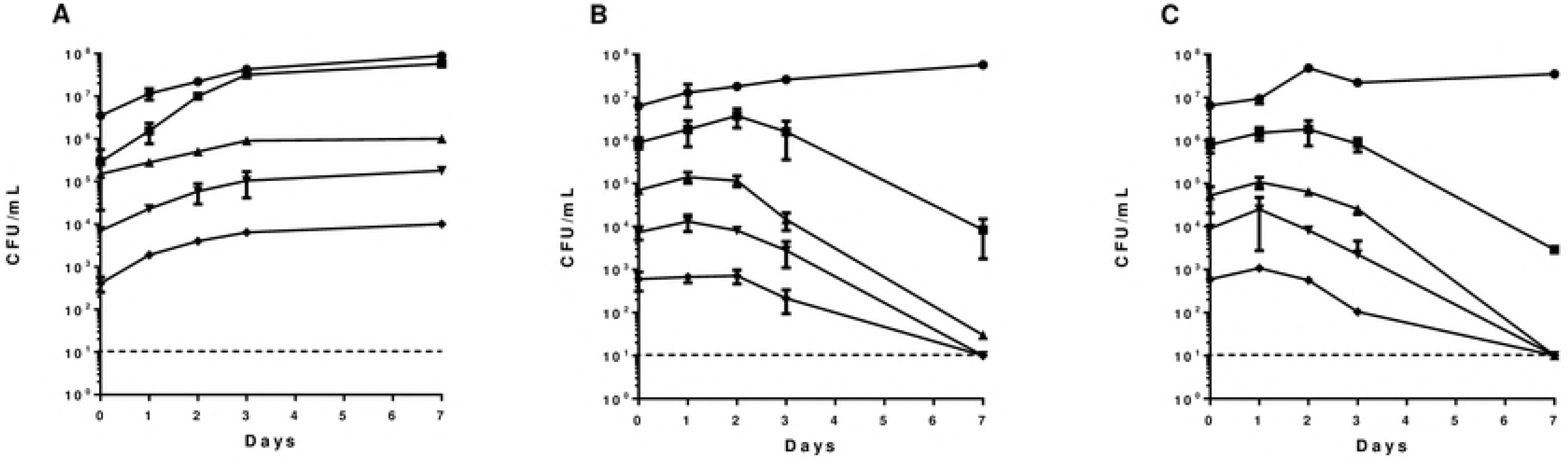
The effects of inoculum size on compound activity. Aerobic cultures were exposed to compounds at 10X MIC over 7 days. Bacterial viability was determined by counting CFU. (A) DMSO, (B) Compound **1**, (C) **2**. The starting inoculum was: (•)10^7^, (▪) 10^6^, (▴) 10^5^, (▾) 10^4^, and (♦) 10^3^.The dashed line represents the lower limit of detection.

### PHY compounds are rapidly bactericidal against starvation--induced, non-replicating *M. tuberculosis*

One of the complications of LORA, is that it requires a period of outgrowth after exposure to compound under hypoxic conditions when the compound is still present. We used the nutrient starvation model, in which loss of bacterial replication is due to complete starvation in order to determine compound efficacy against non-replicating bacteria. In this model bacilli are starved for 2 weeks before compound exposure, and bacterial viability monitored over 21 days. The dilution step remove any compound carryover during plating. We tested four compounds (Fig 4). Interestingly, all the compounds showed much greater activity i.e. more rapid kill and at lower concentrations, than against replicating bacilli. For all compounds, cultures were sterilized by day 14 even at the lowest concentration tested. Even compound **6**, which had weak activity in aerobic culture, was very active under these conditions.

**Fig 4.**
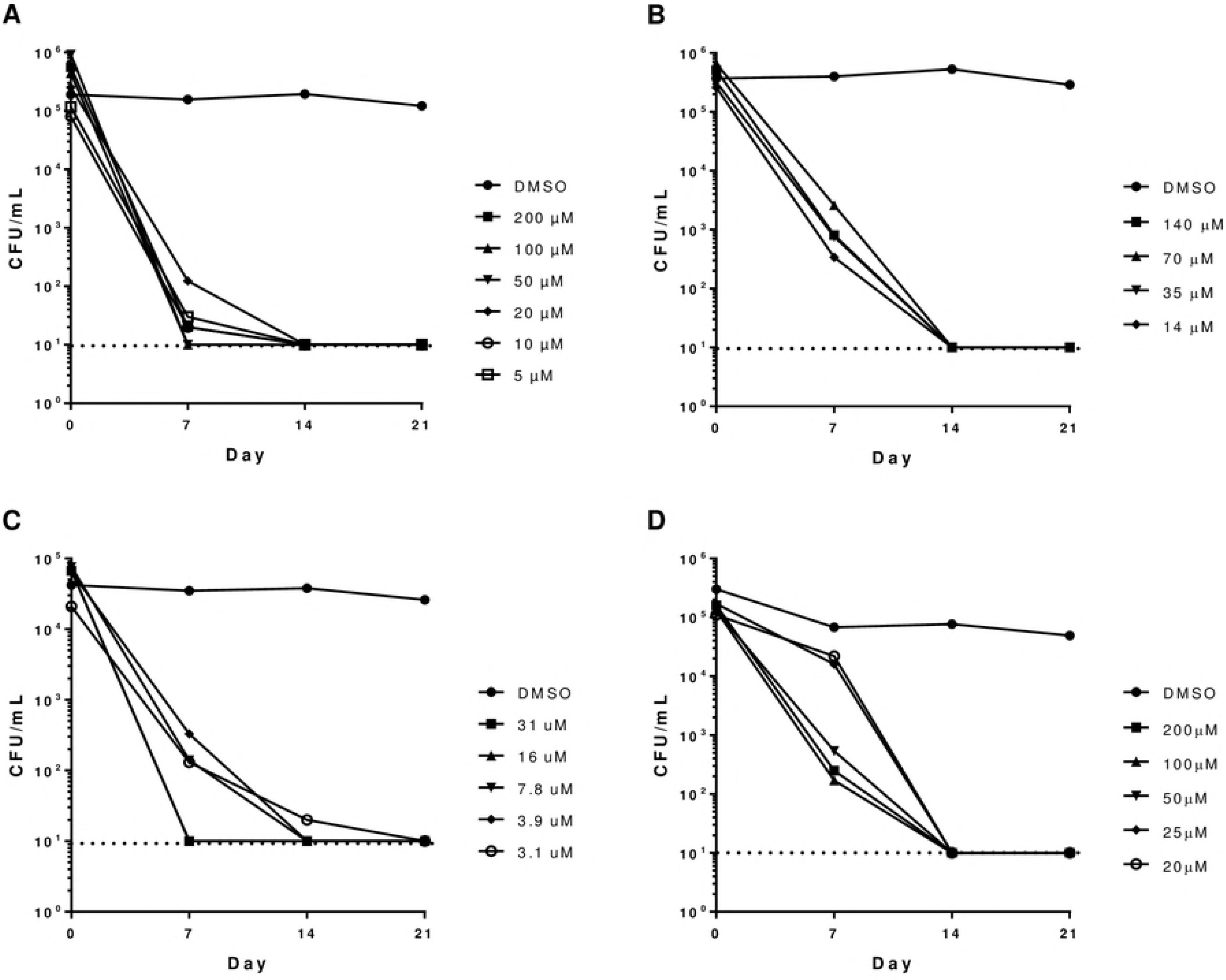
Kill kinetics against non-replicating *M. tuberculosis*. Bacteria were starved in PBS for 2 weeks before compound addition. Bacterial viability in the presence of compound was determined by counting CFU every 7 days over a 21-day period. Compounds (A) **1,** (B) **2**, (C) **3** and (D) **6**. The dashed line represents the lower limit of detection.

### PHY compounds are not active against other bacterial species

We were interested to determine the spectrum of activity of the PHY series. We had previously noted narrow selectivity for *M. tuberculosis* over eukaryotic cells with selectivity indices of <10 [14]. We wanted to determine if this reflected a broad spectrum of activity against all organisms. We measured activity against a range of species on solid medium; for each species we determined the MIC_99_, defined as the minimum concentration required to reduce growth by 99% (Table 2). Activity against *M. tuberculosis* was lower on solid medium than in liquid, with only 2 of the 5 compounds tested showing appreciable activity. The MIC_99_ for compounds **1** and **3** on solid medium was 3.1 and 12.5 μM, respectively, which is 2-6 -fold lower than the MIC in liquid culture. Compounds were tested against representative Gram-negative species (*Escherichia coli* and *Pseudomonas aeruginosa*), as well as Gram-positive (*Bacillus subtilis* and *Staphylococcus aureus*), other mycobacteria (*Mycobacterium smegmatis*) and another eukaryote, *Saccharomyces cerevisiae*. Compounds **1**, **3** were inactive against all species except *M. tuberculosis*. Compounds **4** and **6** had low activity (MIC_99_ = 50-100 μM) against *Sacc. cerevisiae,* with compound **4** showing minimal activity against *B. subtilis* and *Staph. aureus*. There was no correlation between the activity against *M. tuberculosis* and other species, since the most active anti-tubercular compounds were inactive against other species.

**Table 2.** Spectrum of activity for PHY compounds. MIC_99_ were measured on solid medium for each species and defined as the concentration required to inhibit growth by 99%.

| Cpd # | MIC<sub>99</sub> (μM) |  |  |  |  |  |  |
| --- | --- | --- | --- | --- | --- | --- | --- |
|  | *E. coli* | *M. smegmatis* | *S. cerevisiae* | *P. aeruginosa* | *B. subtilis* | *S. aureus* | *M. tuberculosis* |
| 1 | >100 | >100 | >100 | >100 | >100 | >100 | 3.1 |
| 2 | >100 | >100 | >100 | >100 | >100 | >100 | > 100 |
| 3 | >100 | >100 | >100 | >100 | >10 | >100 | 12.5 |
| 4 | >100 | >100 | 50 | >100 | 100 | 100 | > 50 |
| 6 | >100 | >100 | 100 | >100 | >100 | >100 | > 50 |

In conclusion, we have demonstrated that the PHY compounds have some attractive properties in terms of their anti-tubercular activity. *M. tuberculosis* can survive in hostile environments in actively replicating, slow growing or non-replicating states [5, 6, 42, 43]. The presence of slowly replicating and non-replicating persistent forms of *M. tuberculosis* contributes to latent TB infections and drug tolerance, which ultimately leads to the long treatment therapy [12]. Hypoxia and nutrient deprivation are two conditions *M. tuberculosis* encounters during infection. The PHY compounds were active in two single stress models (hypoxia and starvation) under conditions that promote antibiotic tolerance. In addition, they demonstrated higher rates of kill against non-replicating than replicating bacilli. Our previous work had demonstrated that they are also more potent against a strain of *M. tuberculosis* with reduced LepB [14]. Since LepB expression is reduced under both nutrient-starved and hypoxic conditions [6, 7], this may account for the increased activity of PHY compounds in these conditions.

## Acknowledgements

We thank Matthew McNeil for technical assistance.

